# Microtubule-associated protein IQ67 DOMAIN5 regulates interdigitation of leaf pavement cells in *Arabidopsis thaliana*

**DOI:** 10.1101/268466

**Authors:** Dipannita Mitra, Pratibha Kumari, Jakob Quegwer, Sandra Klemm, Birgit Möeller, Yvonne Poeschl, Paul Pflug, Gina Stamm, Steffen Abel, Katharina Bürstenbinder

## Abstract

Plant microtubules form a highly dynamic intracellular network with important roles for regulating cell division, cell proliferation and cell morphology. Its organization and dynamics are coordinated by various microtubule-associated proteins (MAPs) that integrate environmental and developmental stimuli to fine-tune and adjust cytoskeletal arrays. IQ67 DOMAIN (IQD) proteins recently emerged as a class of plant-specific MAPs with largely unknown functions. Here, using a reverse genetics approach, we characterize Arabidopsis IQD5 in terms of its expression domains, subcellular localization and biological functions. We show that *IQD5* is expressed mostly in vegetative tissues, where it localizes to cortical microtubule arrays. Our phenotypic analysis of *iqd5* loss-of-function lines reveals functions of IQD5 in pavement cell (PC) shape morphogenesis, as indicated by reduced interdigitation of neighboring cells in the leaf epidermis of *iqd5* mutants. Histochemical analysis of cell wall composition further suggests reduced rates of cellulose deposition in anticlinal cell walls, which correlate with reduced asymmetric expansion. Lastly, we provide evidence for IQD5-dependent recruitment of calmodulin calcium sensors to cortical microtubule arrays. Our work thus identifies IQD5 as a novel player in PC shape regulation, and, for the first time, links calcium signaling to developmental processes that regulate multi-polar growth in PCs.

**Highlight:** Microtubule-localized Arabidopsis IQ67 DOMAIN5 regulates pavement cell morphogenesis in the leaf epidermis and links calcium-calmodulin signaling to lobe initiation and asymmetric expansion during early phases of interdigitated cell growth.

## Introduction

The plant cytoskeleton, comprised of actin filaments and microtubules (MTs), forms a three-dimensional intracellular network that determines cell division and cell morphology, and serves as tracks for cellular transport of various cargoes, including organelles, proteins and other macromolecular complexes (Akhmanova and Hammer, 2010; Hussey *et al.*, 2006; Wasteneys and Yang, 2004b). Networks of MTs form highly dynamic arrays and adopt specific functions during the plant life cycle. During cell division, MTs organize into the plant-specific preprophase band, which defines the plane of future division (McMichael and Bednarek, 2013), form the mitotic spindle apparatus that facilitates chromosome segregation (Bannigan *et al.*, 2008), and rearrange into the phragmoplast, which serves as scaffold for *de novo* cell plate formation and mediates separation of the two daughter cells (Smertenko *et al.*, 2017). In interphase cells, MTs reorganize into cortical networks tethered to the plasma membrane (PM). Cortical MTs serve as tracks for PM-localized cellulose synthase complexes (CSC) (Liu *et al.*, 2015; Paredez *et al.*, 2006), which assemble long cellulose microfibrils that are the main constituent of plant cell walls. Cellulose fibrils are aligned transversely to the growth axis (Endler and Persson, 2011). The orientation of MTs typically correlates with the direction of cellulose deposition, which collectively determine the direction of cellular expansion during diffuse growth (Gutierrez *et al.*, 2009). In addition to their important roles in development, MT arrays function during growth adaptation in response to changing environmental conditions thereby contributing to plant fitness (Wasteneys and Yang, 2004a).

To engage in these diverse cellular functions, MT organization and dynamics are tightly controlled (Wasteneys, 2002). Developmental and environmental stimuli can induce rapid reorganization of the MT cytoskeleton, e.g. in response to mechanical stimulation, which can occur within a few minutes and involves changes in MT trajectories, as well as altered rates of (de-) polymerization Hardham *et al.*, 2008). Phytohormones exert control over MT orientation (Locascio *et al.*, 2013; Shibaoka, 1994; Takatani *et al.*, 2015), and signaling via the second messenger calcium (Ca^2+^) has been implicated in cytoskeletal control, as suggested by sensitivity of MT stability to elevated Ca^2+^ concentrations (Hepler, 2005, 2016). MT associated proteins (MAPs), which bind to tubulin subunits, play essential roles for regulating cytoskeletal behavior (Lloyd and Hussey, 2001; Sedbrook, 2004) and are likely candidates to integrate incoming signals into appropriate responses. Numerous MAPs have been identified in plants, which mediate bundling, cross-linking, nucleation, or severing of MTs, or, in the case of plus end-tracking MAPs, control dynamic instability at polymerizing plus ends (Akhmanova and Steinmetz, 2008; Horio and Murata, 2014). Other MAPs facilitate physical connections between MTs and protein complexes, such as CSCs (Bringmann *et al.*, 2012), or crosslinking to the actin cytoskeleton (Schneider and Persson, 2015). Moreover, MAPs mediate tethering of MTs to the PM (Bayer *et al.*, 2017; Oda, 2017), which is required for stabilization against the pushing forces of CSCs (Liu *et al.*, 2016) and potentially contributes to sub-compartmentalization of PMs into functional subdomains (Sugiyama *et al.*, 2017). Still, the modes by which external signals are integrated into MT (re-) orientation are poorly understood.

We previously identified IQ67 DOMAIN (IQD) family proteins as the largest known class of MAPs in plants (Bürstenbinder *et al.*, 2017b), which are encoded by multigene families of 23 to 66 members in several angiosperms, including *Arabidopsis thaliana, Oryza sativa* (rice), *Solanum lycopersicum* (tomato), and *Glycine max* (Abel *et al.*, 2005; Feng *et al.*, 2014; Huang *et al.*, 2013). First experimental data point to important roles of IQD proteins in plant development, as indicated by altered fruit shape and grain size in plants with elevated expression levels of tomato *SUN/IQD12* and rice *GRAIN SIZE ON CHROMOSOME 5(GSE5)/IQD21*, respectively (Duan *et al.*, 2017; Xiao *et al.*, 2008). Functions of IQDs in MT organization are supported by differential MT patterns, which are induced upon overexpression of individual family members in transient expression assays in *Nicotiana benthamiana* and transgenic Arabidopsis plants (Bürstenbinder *et al.*, 2017b). The family-defining IQ67 domain harbors motifs with predicted roles in binding to calmodulin (CaM) Ca^2+^ sensor proteins that are integral part of the cellular Ca^2+^ decoding machinery (Abel *et al.*, 2005). Thus, IQDs are likely candidates for integration of CaM-dependent Ca^2+^ signaling into MT (re-)organization and growth regulation (Bürstenbinder *et al.*, 2017a). Mechanistic studies on IQD functions, however, are still limited because (i) multigene families are not easily amenable to reverse genetics approaches due to functional redundancies, and (ii) insights into the spatial and temporal control of Ca^2+^ signal generation during development are sparse due to limited sensitivities of intracellular Ca^2+^ imaging methods (Kudla *et al.*, 2018).

To identify functions of family members, we selected Arabidopsis IQD5, because MT pattern analysis upon overexpression of *YFP-IQD5* suggested unique and specific roles of this family member in MT organization (Bürstenbinder *et al.*, 2017b). In this study, we present a systematic analysis of Arabidopsis *IQD5* using reverse genetics approaches. We identified expression domains of *IQD5* by analysis of transgenic *pIQD5::GFP-GUS* reporter lines, and determined its subcellular localization in transgenic *pIQD5::IQD5-GFP/iqd5-1* lines. We show that IQD5-GFP decorates cortical MTs at neck regions of leaf epidermis pavement cells (PCs). Loss of *IQD5* results in strongly reduced growth restriction at neck regions, which correlates with a reduced deposition of cellulose in anticlinal walls of PCs. Recombinant IQD5 interacts with apo-CaM and Ca^2+^-CaM *in vitro*, and recruits CaM to MTs *in planta*. Together, our research provides evidence for functions of IQD5 in shape establishment of leaf epidermis PCs and points to important roles of Ca^2+^ ions during interdigitated growth of PCs.

## Materials and methods

### Plant material, growth conditions and macroscopic phenotyping

Wild type (WT) seeds (Col-0 accession) were originally obtained from the Arabidopsis Biological Resource Center. T-DNA insertion lines SALK_015580 and GK-288E09, referred to as *iqd5-1* and *iqd5-2*, respectively, were obtained from the Nottingham Arabidopsis Stock Centre. Genomic DNA was extracted as described in Bürstenbinder *et al.* (2007). Homozygous mutant lines were identified by PCR-based genotyping with following primer combinations: *iqd5-1*, WT allele IQD105 (5’-GATTATCTCTGCCAAACAGCG-3’) and IQD106 (5’-GGAGAGTGACTTGGGCTGAC-3’), insert IQD105 + A004 (5’-ATTTTGCCGATTTCGGAAC-3’); *iqd5-2*, WT allele IQD075 (5’-ATGGGAGCTTCAGGGAGATG-3’) + IQD076 (5’-GCGTTACAGCAGCTTGTTTTC-3’), insert IQD076 + A009 (5’-ATAATAACGCTGCGGACATCTACATTT-3’). A 2,201 bp (*pIQD5*_*long*_) and a 1,207 bp (*pIQD5*_*short*_) of the *IQD5* promoter sequence were amplified from genomic DNA with IQD180 (5’-CACCTCTATATATGGTTCACAATCGAGACAC-3’) and IQD345 (5’-CACCATAAATCACATCACTGTTTTTGGGT-3’) forward primers, respectively, in combination with the IQD181 reverse primer (5’-TCTATCTCAATTCCAACGATCAG-3’), and mobilized into the pENTR/dTOPO vector. A genomic *pIQD5_short_::IQD5(w/-stop codon)* fragment was amplified with forward primer IQD1521 (5’-attB1-TCTCTATATATGGTTCACAATCGAGACAC-3’) and reverse primer IQD1522 (5’-attB2-CTGCAAGCCTCTGTTTTATTGGGTCGG-3’), and mobilized into pDONR221. Fidelity of inserts was verified by sequencing. For generation of transgenic *pIQD5::GFP-GUS* and *pIQD5::IQD5-GFP* lines, the inserts were mobilized into pBGWFS7 and pB7FWG,0, respectively (Karimi *et al.*, 2002). Arabidopsis plants were transformed by *Agrobacterium tumefaciens*-mediated transfecting using the floral dip method (Clough and Bent, 1998). Per construct, 10 to 24 independent lines were identified in the T1 generation by Basta selection. T2 plants were screened for the presence of single copy T-DNA insertion by segregation analysis (Basta). For analysis of GFP fluorescence and GUS expression, two to four homozygous T3 lines were included, which showed representative GFP fluorescence or GUS expression patterns.

Seeds were surface sterilized with chlorine gas, stratified for 2 days at 4°C on *Arabidopsis thaliana* Salts (ATS) medium (1x ATS salts, 0.5 % (wv) agar gel, 1 % (w/v) sucrose) (Lincoln *et al.*, 1990), and grown at 21°C under long-day conditions (16 hrs light, 8 hrs dark). For oryzalin treatments, seedlings were incubated for 1-2 hrs in liquid medium supplemented with 10 μM oryzalin in a final concentration of 0.25 % (v/v) DMSO or the DMSO control as described in Bürstenbinder *et al.* (2013). Macroscopic growth parameters were analyzed in 5-day-old seedling and in 3-week-old plants. Root length was quantified with RootDetection (http://www.labutils.de/rd.html). Cotyledon and leaf area were measured with the Easy Leaf Area software (http://www.plant-image-analysis.org/software/easy-leaf-area) according to the manual.

### RNA extraction and expression analysis

Total RNA was extracted from 2-week-old plants using TRIreagent. Synthesis of cDNA via reverse transcription (RT) and RT-PCR reactions were performed according to Bürstenbinder *et al.* (2007) with following primers: *IQD5*, primer IQD075 and IQD117 (5’-CTATGCAAGCCTCTGTTTTATTGG-3’); *ACTIN2*, primers A005 (5’-CAAAGACCAGCTCTTCCATC-3’) and A006 (5’-CTGTGAACGATTCCTGGACCT-3’). For quantitative Real-Time (qRT)-PCR, conditions were selected as described in Bürstenbinder *et al.* (2017b) and following primers were used: *IQD5,* IQD1777 (CAACTAAAGCCAACCGAGCA-3’) and IQD1778 (GGTTTTGGGCAGATTTTTCC-3’), *PP2A* A015 (AGCCAACTAGGACGGATCTGGT-3’) and A016 (CTATCCGAACTTCTGCCTCATTA-3’). In brief, 3 to 5 shoots of 2-week-old plants were pooled for RNA extraction. 2 μg of DNAseI-treated RNA were reverse transcribed with oligo-dT-primers using Revert Aid First Strand cDNA synthesis kit (Thermo Fisher) to generate first-strand cDNA. Primer efficiencies were calculated from standard curves. 1 μl of 1:10-diluted cDNA was used in a 10μl reaction mix including Fast SYBR Green master mix (Applied Biosystems), and qPCRs were run on a 7500 Fast Real-Time PCR system with following program: 10 min, 95 °C; 40 cycles of 3 s, 95 °C and 30 s, 63 °C. Expression levels of *IQD5* were calculated relative to *PP2A*.

### Microscopy, staining procedures and image analysis

Whole mount GUS staining of seedlings and plants was performed as described in Bürstenbinder *et al.* (2017b). Plant materials were cleared in chloral hydrate, and roots and seeds were imaged with a Zeiss axioplan 2 microscope using a DIC objective. Imaging of whole seedlings, leaves, flowers and siliques was performed with a Nikon SMZ 1270 stereo microscope.

Confocal imaging was performed with a Zeiss LSM 780 inverted microscope using a 40x water immersion objective, unless stated otherwise. Generation of YFP-IQD5, Y_N_-TRM1, and mCherry-CaM2 is described in Bürstenbinder *et al.* (2017b) and Gantner *et al.* (2017). GFP was excited using a 488 laser and emission was detected between 493 and 555 nm. YFP was excited by a 514 nm laser, and emission was detected between 525 and 550 nm. For mCherry excitation, a 555 nm laser was used, and emission was detected between 560 and 620 nm. Fluorescence intensities of IQD5-GFP and GFP-MAP4 adjacent to the periclinal wall at convex and concave sides of lobes were quantified according to Armour *et al.* (2015). Average fluorescence intensities were measured with FiJi (Schindelin *et al.*, 2012) in a total of 10 cells from 5 independent seedlings, and 5 lobes per cell were analyzed. The Gehl *et al.* (2009) vector series was used for generation of bimolecular fluorescence complementation (BiFC) construct. For all samples included in the BiFC experiment, imaging was performed with identical laser setting. In co-expression assays, mCherry and GFP fluorescence were recorded in the sequential mode.

For visualization of cell contours, cell outlines were visualized by PI staining as described in Bürstenbinder *et al.* (2017b), and imaged with a 20x objective (5 – 10 days after germination (DAG)) or with a 40x objective (2 and 3 DAG). PI was excited with a 555 nm laser, and emission was detected between 560 and 620 nm. Segmentation, feature quantification, and graphical visualization of PC shapes were conducted with the ImageJ plugin PaCeQuant and the associated R script (Möller *et al.*, 2017). For cells in cotyledons 5 to 10 DAG and in the 3rd true leaf the threshold for size filtering implemented in PaCeQuant was set to the default value of 240 μm^2^. For cells in cotyledons 2 and 3 DAG, the threshold for size filtering was reduced to 75 μm^2^. For time series analysis of cells during cotyledon development, cells were group by their sizes into following categories: tiny, cells ≤ 240 μm^2^; small, 240 − 1400 μm^2^; medium, 1,400 – 4,042 μm^2^; and large, cells ≥ 4,042 μm^2^. Thresholds for small, medium and large cells where chosen according to Möller et al. (2017).

For histochemical cellulose staining in cell walls, 5-day-old seedlings were incubated for 90 min in 0.04 % (v/v) calcofluor-white M2R dissolved in Tris HCl buffer (pH 9.2). To stain callose and the cuticle, seedlings were incubated for 3 hrs and 5 min in 0.1% (v/v) aniline blue in 100 mM Na_2_PO_4_ buffer (pH 7.2) and 0.1% (w/v) auramine O in 50 mM Tris HCl buffer (pH 7.2), respectively. Subsequently, seedlings were co-stained with PI stain to visualize cell contours. Dissected cotyledons were imaged with a Zeiss LSM 700 inverted microscope, using a 40x water immersion objective. Calcofluor-white, aniline blue and auramine O were excited with a 405 nm laser, and emission was detected with a 490 nm short pass filter. Co-staining was recorded in the sequential mode.

To quantify fluorescence intensities along the boundaries of the cells we established a workflow combining automatic segmentation based on the method implemented in PaCeQuant and quantification of fluorescence intensities along the contour segments. For each boundary pixel in an image the set of adjacent cell regions in a 15 × 15 neighborhood around the pixel is determined, and the fluorescence intensity value of the pixel is added to the total intensity sum of each of these regions. Finally, an average intensity value for the boundary of each cell region is calculated by dividing the intensity sum of the region through the total number of pixels that contributed to the specific region, which we implemented in MiToBo (Möller *et al.*, 2016).

### Structure prediction, protein expression and calmodulin binding assays

Structural prediction of the IQ67 domain of IQD5 spanning amino acids E87 to L153 was performed using PHYRE2 (Kelley *et al.*, 2015), which revealed highest similarities with the crystal structures of the CaM binding domains of mouse myosin V (PDB:2IX7, 99.5% similarity) (Houdusse *et al.*, 2006) and mouse myosin-1c (PDB:4R8G, 99.8%) (Lu *et al.*, 2015). The predicted structure of the IQ67 domain of IQD5 was aligned with PDB:2IX7, which contains the crystal structure of apo-CaM bound to the first two IQ motifs of myosin V, using PyMol (DeLano, 2009). CaM was fitted to adjust for the different spacing of IQ-motifs by 11 and 12 amino acids in the CaM binding domains of IQD5 and myosin V, respectively.

Expression of GST-IQD5 and *in vitro* CaM binding assays were performed according to Levy *et al.* (2005). Generation of IQD5 pENTR vectors is described in Bürstenbinder *et al.* (2017b). The CDS of IQD5 was mobilized into the pDEST15 vector (Invitrogen) to generate an N-terminal GST fusion, and GST-IQD5 and the GST control were expressed in the *Escherichia coli* strain KRX (Novagen) upon induction with 0.1% (w/v) rhamnose and 1 mM IPTG. Cells were resuspended in CaM pull-down buffer (5.8 mM Tris-HCl, pH 7.3; 2.7 mM KCl; 127 mM NaCl; 0.1% (v/v) Tween 20; 0.002% (w/v) NaN_3_). Bovine CaM immobilized on sepharose beads (GE Healthcare) was incubated with cleared protein extracts in CaM buffer containing either 5 mM EGTA or 1 mM CaCl_2_. After four steps of washing, the last washing fraction and the bead fraction were collected, and, together with the unbound fraction, separated by SDS-PAGE. GST-tagged proteins were visualized by immuno-blot analysis using a HRP-coupled α-GST antibody (Santa Cruz).

### Statistical analysis

Statistical analysis of root length, cotyledon and leaf area, and *IQD5* expression was performed using ANOVA implemented in the R software, followed by a Tukey’s posthoc test, and Benjamini Hochberg adjustment of p values. For statistical analysis of fluorescence intensities at convex and concave sides of pavement cells in *pIQD5::IQD5-GFP* seedlings, a t-test was performed. Statistical analysis of PC shape features was performed using the Kruskal-Wallis test, followed by a Dunn’s posthoc test and Benjamini-Hochberg-adjustment of p values, which is part of the R script provided in the PaCeQuant package.

## Results

### *IQD5* is expressed in vegetative tissues

To identify *in planta* sites of IQD5 function, we determined spatio-temporal expression domains of *IQD5* in transgenic *pIQD5*_*short*_*::GFP-GUS* and *pIQD5*_*long*_*::GFP-GUS* reporter lines, in which a 1,207 bp and 2,201 bp DNA fragment upstream of the translational start site of the *IQD5* gene were fused to the reporter, respectively. Histochemical GUS analysis of *pIQD5*_*short*_*::GFP-GUS* lines throughout development revealed strong promoter activity in cotyledons and leaves, in the vasculature of leaves and the hypocotyl, as well as in the shoot apical meristem (Fig. 1A). In roots, GUS staining was detectable mostly in older parts of the root. In root tips, *IQD5* promoter activity was restricted to the lateral root cap of primary and lateral root meristems. GUS activity was largely absent from reproductive organs, such as flower buds, flowers, siliques and seeds, and during embryo development. The GUS patterns are consistent with developmental *IQD5* expression data obtained from publicly available microarray datasets (Fig. 1B) (Winter *et al.*, 2007), which confirm higher *IQD5* expression levels in vegetative tissues when compared to reproductive tissues. Similar expression patterns were observed in *pIQD5*_*long*_*::GFP-GUS* lines (Fig. S1), suggesting that the 1,207 bp fragment was sufficient to report authentic *IQD5* expression patterns. Our analysis thus reveals preferential expression of *IQD5* in vegetative tissues of shoots and roots.

**Fig. 1.**
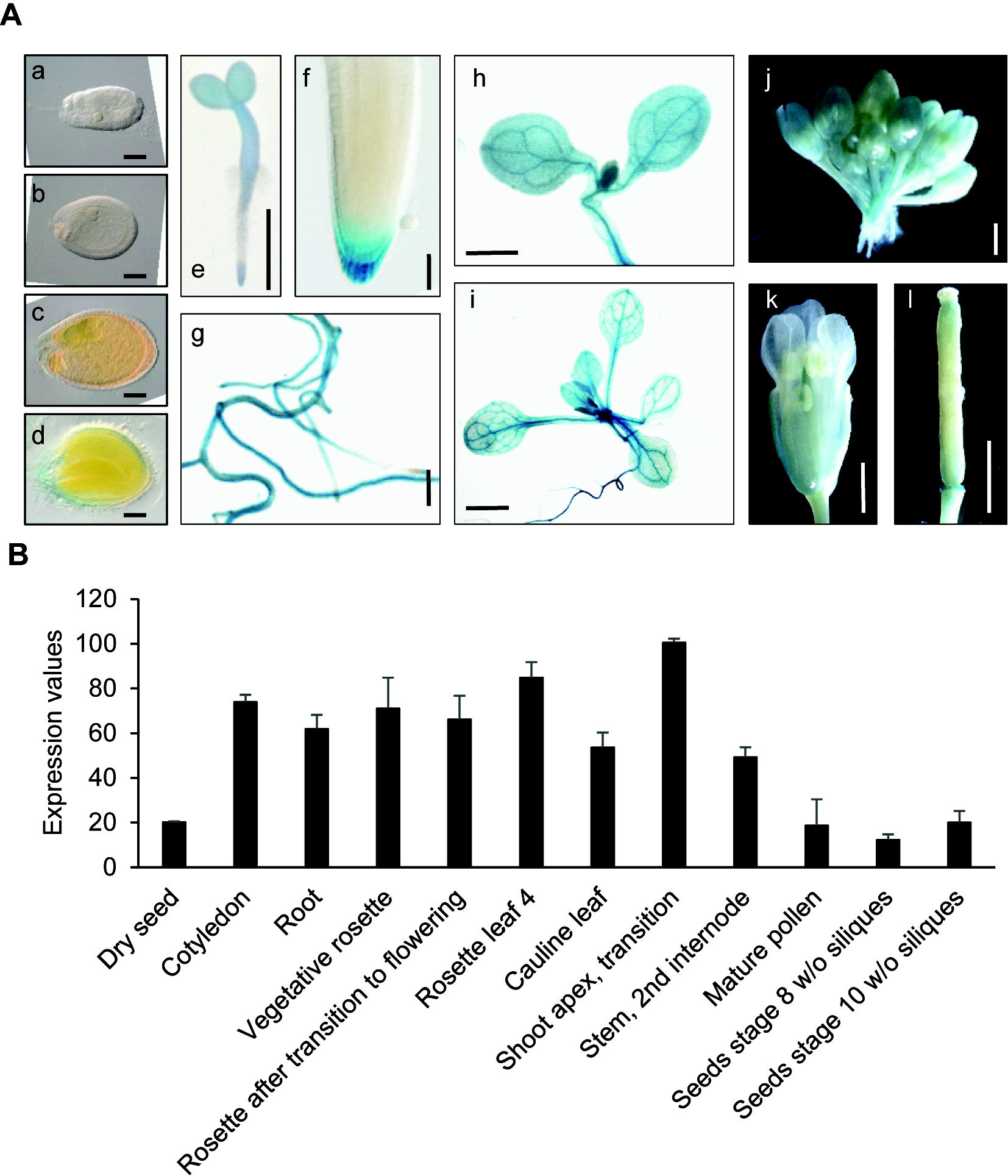
*IQD5* expression analysis. Whole mount histochemical GUS staining of *ProIQD5*_*short*_*:GFP-GUS* reporter lines (A) in seeds and embryos of globular (a), heart (b), torpedo (c) and mature (d) stage, in 2-day-old seedlings (e), in the primary root meristem (f), in lateral roots (g) and cotyledons (h) of 5-day-old seedlings, in the shoot of 10-day-old seedlings (i), and in flower buds (j), flowers (k) and siliques (l) of 5-week-old plants. Bars represent 100 μm (a-d); 1 mm (e, g-l); 10 μm (f). *In silico* expression data of *IQD5* in different tissues and organs were obtained from the publicly available eFP browser database (http://bar.utoronto.ca/efp/cgi-bin/efpWeb.cgi) (B). Data show mean values ± standard deviation from three independent biological experiments.

### IQD5-GFP localizes to cortical microtubules

To examine the subcellular localization of IQD5, we generated a fluorescent protein fusion construct, in which GFP was fused to the C-terminus of IQD5 within a genomic fragment containing the native *IQD5*_*short*_ promoter (Fig. 2). The *pIQD5::IQD5-GFP* construct was introduced into an *iqd5* knockout background to avoid dosage-effects of *IQD5* copy-number (Fig. 2A and 2B). We obtained two independent Arabidopsis T-DNA insertion lines for *IQD5*, which we termed *iqd5-1* and *iqd5-2* (Fig. 2A). RT-PCR analysis revealed complete absence of full-length *IQD5* transcripts in *iqd5-1* and *iqd5-2* lines when compared to the WT, demonstrating that both T-DNA insertion lines are null mutant alleles (Fig. 2B). Based on macroscopic examination, both *iqd5* mutants were phenotypically indistinguishable from WT plants, as shown for root length and shoot growth (Fig. S2). The *iqd5-1* mutant was transformed with the *pIQD5::IQD5-GFP* construct by *Agrobacterium*-mediated floral dip. Quantitative RT-PCR analysis of steady-state *IQD5* mRNA levels revealed comparable expression in two independent *pIQD5::IQD5-GFP/iqd5-1* complementation lines, which was moderately higher than in the reference WT (Fig. 2C).

**Fig. 2.**
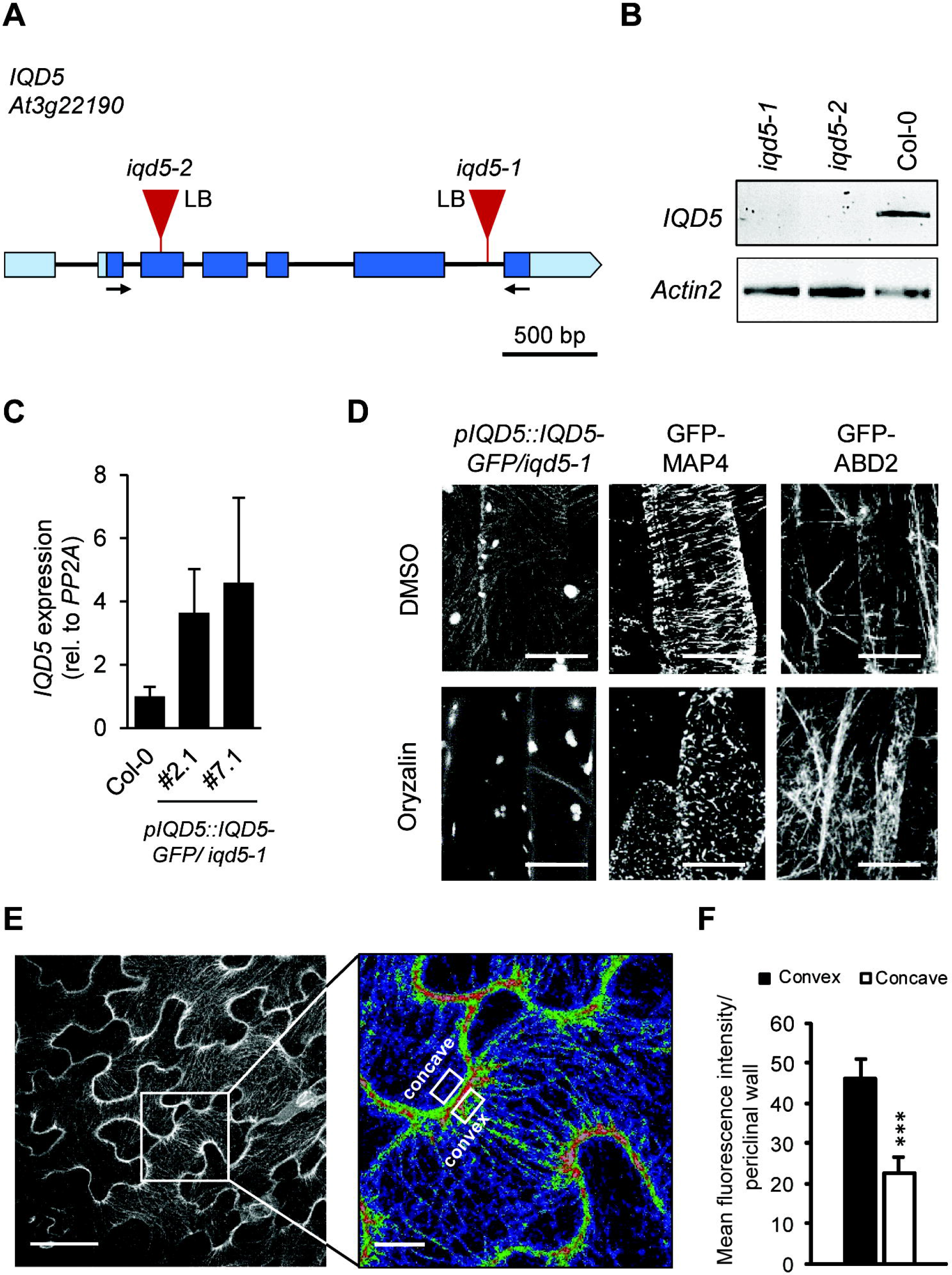
Subcellular localization of IQD5-GFP in transgenic Arabidopsis *pIQD5::IQD5-GFP/iqd5-1* lines. Gene model and position of T-DNA insertions in two independent mutant lines, *iqd5-1* and *iqd5-2* (A). Boxes indicate 5‘UTR and 3‘UTR (light blue) and exons (dark blue). Introns are represented by the black line. Loss of *IQD5* full-length transcript in *iqd5-1* and *iqd5-2* plants compared to the WT (Col-0) was validated by RT-PCR (B). Arrows in (A) indicate the position of primers used for amplification of *IQD5* transcripts. *Actin2* was included as control for cDNA integrity. Relative *IQD5* expression levels were analyzed by qRT-PCR in two independent transgenic *pIQD5::IQD5-GFP/iqd5-1* lines (#2.1 and #7.1) compared to the WT (C). Data show mean values ± standard deviation of three independent biological experiments. Subcellular localization of IQD5-GFP in hypocotyls of transgenic *pIQD5::IQD5-GFP/iqd5-1* seedlings after mock (DMSO) or oryzalin (10 μM) treatment; bars, 10 μm (D). Transgenic *pCaMV35S::GFP-MAP4* and *pCaMV35S::GFP-ABD2* seedlings were included as controls for the microtubule and actin cytoskeleton, respectively. Z-stack images of GFP fluorescence in epidermis pavement cells of cotyledons from 5-day-old *pIQD5::IQD5-GFP/iqd5-1* seedlings (E). Overview images (left column) and close-up of lobe regions (right column, fluorescence intensities are shown by false coloring, red, high fluorescence intensity; blue, low fluorescence intensity). Bars, 50 μm and 5 μm, respectively. Mean fluorescence intensities measured at the convex and concave site of lobe regions in the upper periclinal wall (F). Data show mean values ± standard deviation from a total of 50 lobes, quantified in 10 cells from 5 seedlings.

We investigated IQD5-GFP subcellular localization by confocal imaging and observed that IQD5 localized in punctate patterns along filamentous structures at the cell cortex of hypocotyl cells, reminiscent of cortical MTs (Fig. 2D). Treatment with oryzalin a drug that binds to tubulin subunits and prevents MT polymerization (Morejohn *et al.*, 1987), abolished IQD5-GFP localization to filaments, while MTs remained intact upon mock treatment (Fig. 2D). Transgenic *pCaMV 35S::GFP-MAP4* (Marc *et al.*, 1998) and *pCaMV35S::GFP-ABD2* (Sheahan *et al.*, 2004; Wang *et al.*, 2004) lines were included as controls for the MT and actin cytoskeleton, respectively (Fig. 2D). While oryzalin treatment disrupted MTs decorated with GFP-MAP4, labeled (GFP-ABD2) actin filaments remained intact, demonstrating efficiency and specificity of the treatment.

GFP fluorescence was very weak in hypocotyls of *pIQD5::IQD5-GFP/iqd5-1* seedlings. Moderately stronger IQD5-GFP fluorescence was detectable at cortical MT arrays in epidermal PCs of cotyledons (Fig. 2E). PCs adopt highly complex jigsaw puzzle-like shapes with interlocking lobes and necks. Within individual PCs, IQD5-GFP accumulated at the convex side of necks at the interface of anticlinal and outer periclinal walls, as indicated by increased average fluorescence intensities when compared to the concave side (Fig. 2E and 2F). Similarly, cortical MTs accumulate at convex sides of necks during PC development, and growth is largely restricted at these neck regions (Armour *et al.*, 2015).

### Loss of *IQD5* causes aberrant PC shape

To assess whether IQD5 contributes to growth regulation of PCs, we analyzed PC shapes in cotyledons of *iqd5* mutants and of the two *pIQD5::IQD5-GFP/iqd5-1* lines. Cell outlines were visualized by PI staining, and groups of PCs on the adaxial side of cotyledons from 5-day-old seedlings were imaged by confocal microscopy (Fig. 3A). Quantification of PC shape features with PaCeQuant (Möller *et al.*, 2017), an ImageJ-based open source tool that we developed for fully automatic quantification and graphical visualization of PC shape features, revealed that cell expansion was largely unaffected in *iqd5* mutants, as evidenced by similar sizes of individual cells and similar size distributions (Fig. 3B). Cell shapes on the other hand differed strongly in both *iqd5* mutant alleles when compared to WT or the two independent *pIQD5::IQD5-GFP/iqd5-1* lines. The average number of lobes was moderately reduced from 15 lobes per cell in WT to 13 lobes per cell in *iqd5* mutants (Fig. 2C). In addition, *iqd5* mutants displayed a strongly reduced growth of lobes, indicated by an ~30% reduction of average lobe length (Fig. 2D). The width of the cellular core region, measured as minimum (Fig. 3E) and maximum (Fig. S3A) core width, were increased by 35% and 22%, respectively. The reduced lobe growth and more uniform expansion of individual PCs is reflected by increased cellular circularity (Fig. 2F) and solidity (Fig. S3A), and a less irregular cell contour as reflected by a reduced margin roughness (Fig. S3A). The phenotypic differences were highly similar between *iqd5-1* and *iqd5-2* mutant alleles, further supporting complete *IQD5* gene knockout in both lines. Expression of *pIQD5::IQD5-GFP* in the *iqd5-1* mutant background restored PC shape to WT-like patterns (Fig. 3, Fig. S3), which demonstrates functionality of the IQD5-GFP fusion protein and sufficiency of the amplified promoter region for restoring *IQD5* expression levels. Collectively, our data suggest that IQD5 is required to restrict growth at neck regions of cotyledon PCs, which is consistent with its predominant localization to cortical MT arrays at necks.

**Fig. 3.**
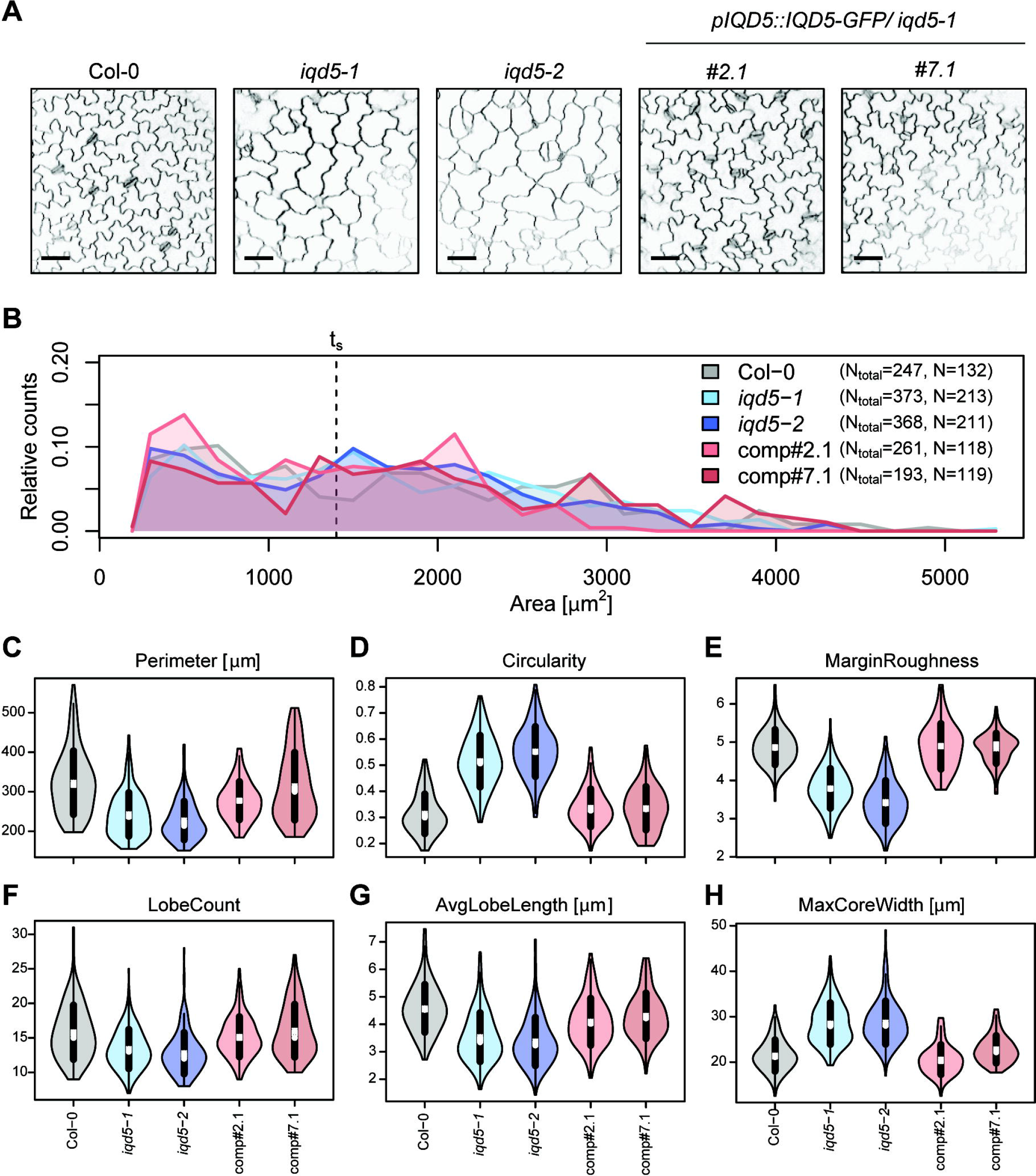
Pavement cell (PC) morphology on the adaxial side of cotyledons from 5 day-old seedlings of wild-type (Col-0), two independent *iqd5* knock-out lines (*iqd5-1* and *iqd5-2)* and two independent transgenic complementation lines (*pIQD5:IQD5-GFP/iqd5-1;* lines *#2.1 and #7.1)*. Representative images of PC morphology (A). Cell outlines were visualized with PI. Images are single optical sections. Bars, 50 μm. Quantification of cell shape features by PaCeQuant (B-F). Relative distribution of cell areas in the analyzed genotypes (B). Numbers in the legend refer to the total number of cells from 10 images of the different genotypes. Cells larger than size threshold t_s_ = 1400μm^2^ were used for further analysis. Violin plots of feature distributions for circularity (C), lobe count (D), average lobe length (AvgLobeLength, E), and maximum core width (MaxCoreWidth, F). Circles and crosses refer to medians and means, the vertical black lines represent standard deviation (thick lines) and the 95% confidence intervals (thin lines). The width of each violin box represents the local distribution of feature values along the y axes. For an overview of all shape features and statistical analysis see Supplemental Fig. S3.

### Cell shape defects in *iqd5* mutants occur early during cotyledon development

Morphogenesis of PCs in cotyledons is established during distinct phases (Fu *et al.*, 2002; Zhang *et al.*, 2011). In the early phase, 1-3 days after germination (DAG), lobe formation is initiated and cells start to expand asymmetrically, which is followed (3-7 DAG) by diffuse growth and expansion of shape patterns. Growth ceases at later stages (10-18 DAG), and PCs as well as cotyledons reach their final size (Belteton *et al.*, 2018). To determine at which stage IQD5 functions, we studied PC shape in *iqd5* mutants during cotyledon development and imaged WT and *iqd5* mutant seedlings at 2, 3, 5, 7 and 10 DAG (Fig. 4A). During development, the average cell size increased (Fig. S4-S8), which is consistent with earlier reports (Möller *et al.*, 2017; Zhang *et al.*, 2011). We applied the size thresholds of t_s_ = 1,400 μm^2^ and t_m_ of 4,040 μm^2^, which we experimentally determined in our previous work (Möller *et al.*, 2017), to examine shape and geometries in cell populations of similar sizes, referred to as small, medium and large, at 3, 5, 7 and 10 DAG. To distinguish between very small (tiny) and small cell populations in cotyledons at 2 and 3 DAG, we included an additional size threshold of t_tiny_ = 240 μm^2^. Quantification of PC shape features revealed first differences in cell shapes of *iqd5* mutants already at 2 and 3 DAG (Fig. 4C, D). When compared to WT, cellular circularity was moderately but significantly increased in both *iqd5* mutant alleles in tiny and in small-sized cell populations (Fig. 4C), and margin roughness as well as average basal lobe length were reduced (Fig. 4D, Fig. S4). Similar results were observed in seedlings at 3 DAG (Fig. 4 E, F, Fig. S5). In medium to large-sized cell populations, analyzed in cotyledons between 5 and 10 DAG, phenotypic differences became more pronounced with increasing cell size (Fig. 4G, Fig. S6-S8). The time series analysis thus suggests important roles of IQD5 already during early phases of PC morphogenesis in cotyledons. Analysis of *pIQD5*_*short*_*::GFP-GUS* revealed promoter activity in cotyledons and in the shoot apical meristem between 2 and 10 DAG (Fig. 4B). Thus, our data demonstrate that *IQD5* is expressed early during cotyledon development and that loss of *IQD5* causes reduced lobe initiation and asymmetric expansion during early growth phases.

**Fig. 4.**
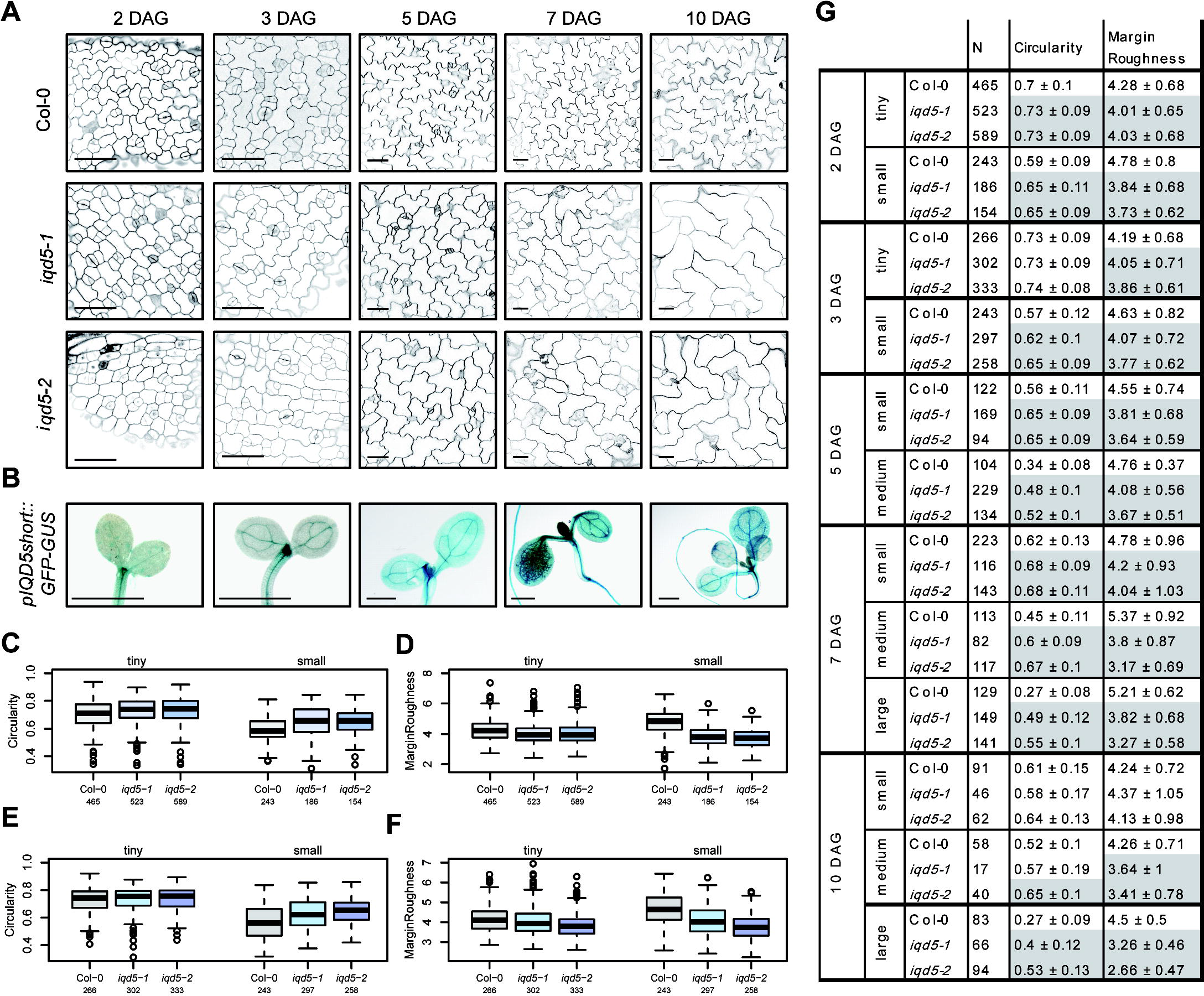
PC morphology during cotyledon development. Representative images of epidermis cells on the adaxial side of cotyledons of WT, *iqd5-1* and *iqd5-2* seedlings at 2, 3, 5, 7, and 10 days after germination (DAG) (A). Cell outlines were visualized by PI; bars, 50 μm. Histochemical GUS staining of *pIQD5*_*short*_*::GFP-GUS* seedlings at indicated time points; bars, 1 mm (B). Quantification of PC shape parameters by PaCeQuant. Cell populations were grouped according to their size as tiny, 75 – 240 μm^2^; small, 240 – 1400 μm^2^; medium, 1400 – 4042 μm^2^; large ≥ 4042 μm^2^. Boxplots show feature distributions for circularity (C, E) and margin roughness (D, F) in seedlings at 2 DAG (C, D) and at 3 DAG (E, F). Results are medians, boxes range from first to third quartile. Feature values for circularity and margin roughness in tiny, small, medium and large-sized cell population during cotyledon development (G). Results show mean values ± standard deviation. Statistically significant differences (p ≤ 0.05) of *iqd5* mutants relative to WT are highlighted in gray. For an overview of all shape features and statistical analysis see Supplemental Fig. S4 – S8.

### IQD5 regulates pavement cell shape during embryogenesis and post-embryonic growth

Cotyledons resemble true leaves in many aspects and thus provide a convenient system to study leaf development (Tsukaya *et al.*, 1994). However, while cotyledons emerge in embryogenesis, true leaves post-embryonically differentiate from the shoot apical meristem and, unlike cotyledons, differ in their final leaf shape (Tsukaya, 2002). Moreover, some mutations affect exclusively the development of cotyledons or true leaves (Tsukaya, 1995). To test if *IQD5* also functions in true leaf development, we analyzed PC shape in rosette leaves of 3-week-old plants (Fig. 5A). Morphologically, the first two true leaves in Arabidopsis are similar to cotyledons (Kerstetter and Poethig, 1998; Poethig, 1997), and phenotypes in some rosette leaf-specific mutants are only visible beyond the second true leaf (Guo *et al.*, 2015). To reflect characteristics of true leaves, we thus focused on the third rosette leaf and analyzed PC shape on the adaxial side (Fig. 5C). Quantification of PC shape features revealed similar shape defects in rosette leaves as observed for cotyledons. Loss of *IQD5* caused a reduced initiation of lobes, as indicated by reduced lobe counts in *iqd5-1* and *iqd5-2* mutants when compared to WT (Fig. 5F), and the average lobe length of *iqd5* mutants was reduced (Fig. 5G). Reduced formation and growth of lobes was additionally reflected by increased circularity (Fig. 5D) and solidity (Fig. S9A) values, as well as reduced margin roughness in PCs of *iqd5* mutant plants. Values of minimum (Fig. S9A) and maximum core width (Fig. 5H) increased, indicative of reduced growth restriction at neck regions. IQD5 thus controls lobe initiation and asymmetric expansion in cotyledons and true leaves. In agreement with functions of IQD5 in true leaves, histochemical GUS activity was detectable in *pIQD5*_*short*_*::GFP-GUS* lines within the entire leaf, indicating that *IQD5* is expressed throughout PC growth (Fig. 5B). Taken together, our data identify IQD5 as a novel regulator of leaf epidermis PC shape, which controls growth restriction at necks in embryonic and post-embryonic tissues.

**Fig. 5.**
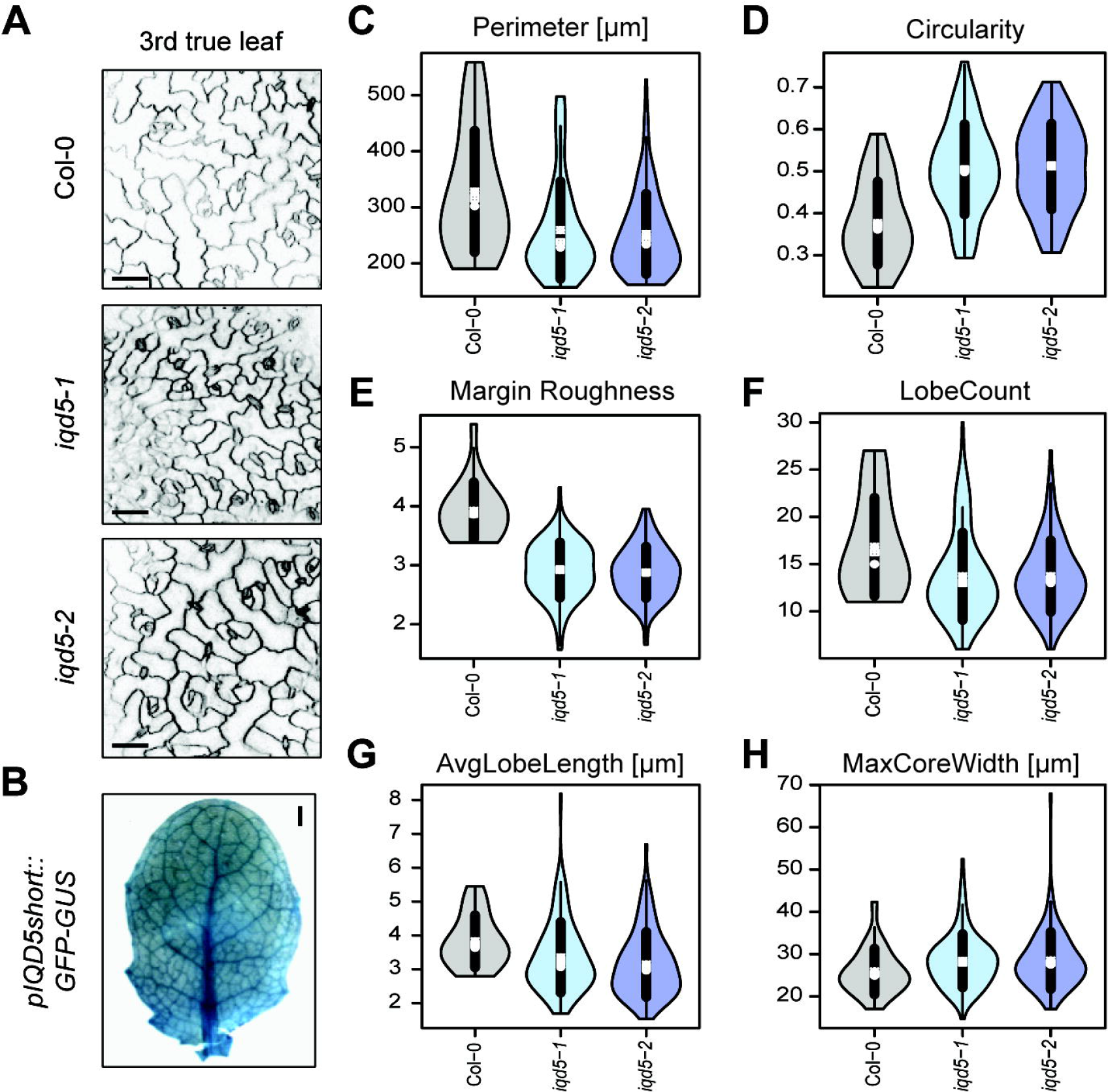
PC shapes in the epidermis of rosette leaves of WT and *iqd5* mutants. Images are single optical sections of PI-labeled epidermis cells on the adaxial side of the third rosette leaf in 3-week-old plants; bars, 50 μm (A). Whole-mount GUS staining of the third rosette leaf in 3-week-old *pIQD5*_*short*_*::GFP-GUS* plants; bar, 1 mm (B). Quantification of PC shape features. Violin plots show feature distributions from n = 23 − 110 cells from 9 − 13 images for perimeter (C), circularity (D), margin roughness (E), lobe count (F), average lobe length (G), and maximum core width (H). Circles and crosses refer to medians and means, the vertical black lines represent standard deviation (thick lines) and the 95% confidence intervals (thin lines). The width of each violin box represents the local distribution of feature values along the y axes. For an overview of all shape features and statistical analysis see Supplemental Fig. S9.

### Reduced growth restriction correlates with altered cellulose deposition

MTs guide cellulose synthase complexes and determine the deposition and direction of newly forming cellulose fibrils in the cell wall (Endler and Persson, 2011; Gutierrez *et al.*, 2009; Paredez *et al.*, 2006). In PCs, cellulose microfibrils accumulate in neck regions at the interface of anticlinal and outer periclinal walls (Panteris *et al.*, 1993), which correlates with sites of MT bundling and local restriction of cell expansion (Armour *et al.*, 2015; Qiu *et al.*, 2002). Because IQD5-GFP preferentially accumulates in neck regions and mutants defective in *iqd5* display shape defects reminiscent of decreased growth restriction at necks, we aimed to investigate whether IQD5 affects cellulose deposition. Staining with calcofluor white, a dye used for visualization of cellulose fibrils (Anderson *et al.*, 2010; Seagull, 1986), revealed reduced staining intensities in *iqd5-1* and *iqd5-2* when compared to WT, which were reverted to WT levels in the complementation line (Fig. 6A). To quantitatively assess differences in fluorescence intensities at anticlinal cell walls, we segmented the contour of individual cells after visualization of cell walls by co-staining with PI using the segmentation mode implemented in PaCeQuant. We measured a ~45% reduction of calcofluor white fluorescence intensities along the cell contour of *iqd5* mutant cells compared to WT and the complementation line (Fig. 6E). Reduced intensities suggest reduced deposition of cellulose in anticlinal cell walls of PCs. Calcofluor white, however, does not discriminate between β-1,3 and β-1,4-glucan chains (Anderson *et al.*, 2010), which are the building blocks of callose and cellulose, respectively. To test whether lesions in *iqd5* specifically affect cellulose deposition, we included aniline blue staining to visualize callose (Wood, 1984) and quantified fluorescence intensities. No differences in callose deposition were observed in the mutants when compared to the WT or the complementation lines (Fig. 6B, E). Similarly, only minor differences (5-10%) in fluorescence intensities were observed upon auramine O staining, which labels the cuticle (Considine and Knox, 1979). Taken together, our data suggest that the reduced calcofluor white signals are not an artefact of reduced penetration or uptake of the dyes due to general defects in cell wall composition, and likely reflect reduced cellulose deposition caused by the loss of *IQD5*.

**Fig. 6.**
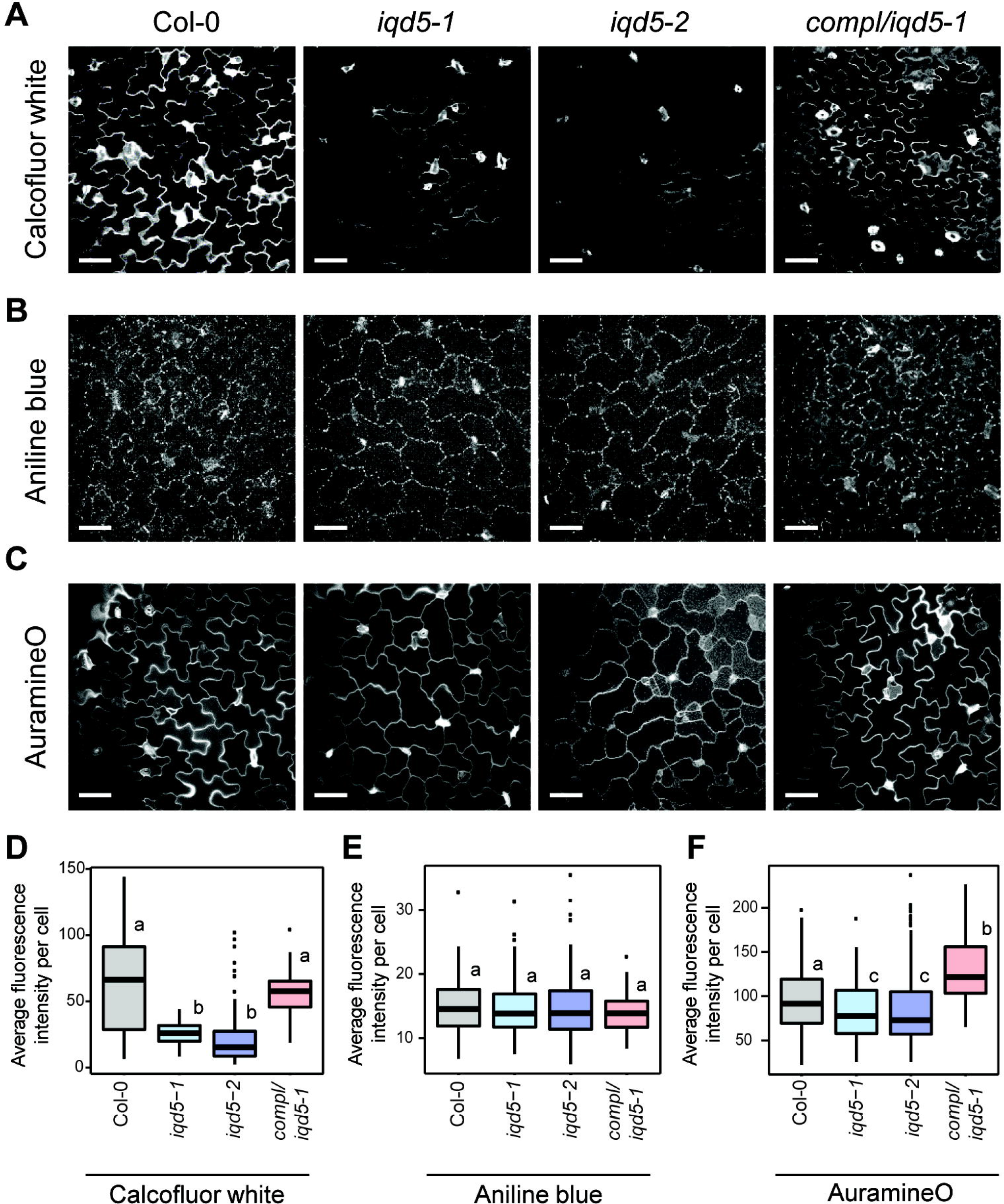
Histochemical analysis of cell wall composition in the WT, the two *iqd5* mutant alleles *iqd5-1* and *iqd5-2*, and in one transgenic *pIQD5::IQD5-GFP/iqd5-1* line. Confocal images show single optical sections of epidermis pavement cells in cotyledons of 5-day-old seedlings. Cellulose staining by calcofluor-white (A), aniline blue staining of callose (B), and auramine O staining of the cuticle (C). Bars, 50 μm. Quantification of relative fluorescence intensities along the anticlinal cell wall (D). Shown are medians, boxes range from first to third quartile. Different letters denote a significant statistical difference by one-way ANOVA, p < 0.01.

### IQD5-dependent recruitment of CaM to cortical microtubules

A hallmark of IQD proteins is the presence of their eponymous IQ67 domain, which contains a repetitive arrangement of predicted CaM binding motifs (Abel *et al.*, 2005). In IQD5, the IQ67 domain contains three copies each of three different classes of CaM interacting motifs, including the IQ motif and motifs of the 1-5-10 and 1-8-14 classes, with presumed roles for binding to apo-CaM and holo-CaM, respectively (Fig. 7A). Homology modeling of IQD5 indicates that the IQ67 domain adopts a α-helical fold (Fig. 7B), similar to the CaM binding domain of myosin (Houdusse *et al.*, 2006), and potentially interacts simultaneously with more than one CaM polypeptide. To assess whether IQD5 is a functional CaM target, we performed *in vitro* CaM binding assays. We expressed GST-tagged IQD5 and the GST core as a control in *E. coli* to investigate interaction with immobilized bovine CaM in the presence (Ca^2+^) and absence (EGTA) of calcium. GST-IQD5, but not GST, co-sedimented with both states of CaM, which suggests that CaM binding is independent of the GST tag, and that the predicted CaM binding motifs are functional (Fig. 7C). To gain insight into subcellular sites of IQD5 interaction with CaM, we performed BiFC analyses. N-terminal fusions of IQD5 to the N-terminal half of YFP (Y_N_-IQD5) were transiently co-expressed with N-terminal fusions of CaM2 to the C-terminal half of YFP (Y_C_-CaM2) in *N. benthamiana* leaves by infiltration with *Agrobacterium* harboring respective plasmids. As controls, we included Y_N_- and Y_C_- fusions of TON1 RECRUITMENT MOTIF1 (TRM1), a member of a plant-specific class of MAPs that interacts with TONNEAU1 (TON1) *in planta* (Drevensek *et al.*, 2012). Recovery of YFP fluorescence was visible along the MT lattice between Y_N_-IQD5 and Y_C_-CaM2, and between Y_N_-TRM1 and Y_C_-TON1, which served as positive control (Fig. 7D). No fluorescence complementation was detectable in the negative controls, in which Y_N_-IQD5 and Y_C_-CaM2 were combined with Y_C_-TRM1 and Y_N_-TRM1, respectively, demonstrating specificity of the BiFC assay. Additionally, CaM binding at MTs was validated in co-expression assays. Expression of *pCaMV35S::mCherry-CaM2* resulted in cytosolic accumulation of mCherry-CaM2, consistent with previous reports (Bürstenbinder *et al.*, 2013). Upon co-expression with YFP-IQD5, mCherry -CaM2 re-localized to cortical MTs (Fig. 7E). Thus, our data point to roles of IQD5 in CaM-recruitment to cortical MTs, and provide first indications for CaM-dependent Ca^2+^ signaling in shape development of leaf epidermis PCs (Fig. 7F).

**Fig. 7.**
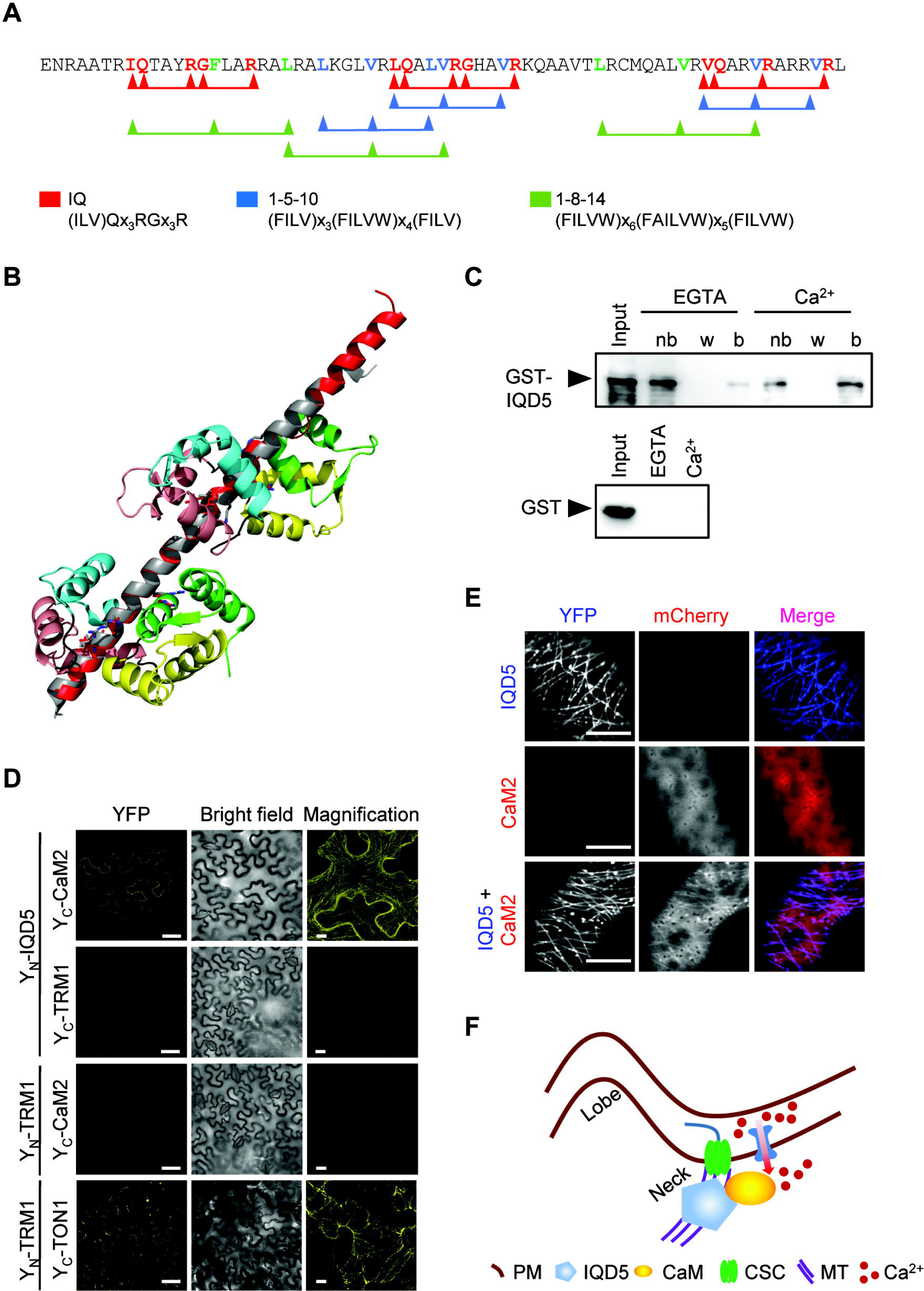
Calmodulin (CaM) binding properties of IQD5. Amino acid sequence of the IQ67 domain of IQD5 (A). IQ motifs, 1-5-10 and 1-8-14 motifs implicated in apoCaM (IQ) and Ca^2+^-CaM (1-5-10 and 1-8-14) binding are highlighted in red, blue and green, respectively. Structural alignment of the IQ67 domain of IQD5 (red) and the IQ motif containing domain of myosin (grey), together with two CaM proteins (1st, 2nd, 3rd, and 4th EF hand in green, yellow, salmon, and cyan, respectively (aligned and fitted with PyMol) (B). *In vitro* pull-down of recombinant GST-IQD5 and as control of GST alone expressed in *E. coli* with bovine CaM immobilized on agarose beads in the presence (Ca^2+^) and absence (EGTA) of calcium; nb, not bound; w, last wash; b, bead-immobilized fraction (C). *In planta* interaction of IQD5 with CaM (D, E). BiFC assays between Y_N_-IQD5 and Y_C_-CaM2 in leaves of *N. benthamiana* (D). Combinations of Y_N_-IQD5 with Y_C_-TRM1 and Y_C_-CaM2 with Y_N_-TRM1 served as negative controls. Y_N_-TRM1 and Y_C_-TON1 were included as positive control. Single optical sections of YFP fluorescence (left column) and corresponding bright field images (center column). Scale bars, 50 μm. Right column, close-up Z-stack images of YFP fluorescence; bars, 10 μm. Subcellular localization of mCherry-CaM2 (top), YFP-IQD5 (middle), and of YFP-IQD5 and mCherry-CaM2 (bottom) in transient (Co)expression assays in leaves of *N. benthamiana* (E). Bars, 5 μm. Proposed model of IQD5 function in pavement cell morphogenesis (F). IQD5 localizes to microtubules and is required for growth restriction at neck regions, possibly by affecting cellulose deposition. Interaction of IQD5 with CaM at microtubules points to important roles of Ca^2+^ signaling during shape establishment.

## Discussion

PCs are the most abundant cell type in the leaf epidermis, which are characterized by their jigsaw puzzle-like shape in Arabidopsis and in several other plant species (Ivakov and Persson, 2013; Jacques *et al.*, 2014). Genetic and pharmacological studies have identified key signaling events during PC morphogenesis that include antagonistic functions of auxin and cytokinin in activation of distinct Rho-like GTPases from plants (ROPs) in lobes and necks (Fu *et al.*, 2005). ROP signaling regulates the actin and MT cytoskeleton via different effector proteins (Fu *et al.*, 2002; Fu *et al.*, 2009), thereby affecting the composition and mechanical properties of the cell wall, eventually resulting in asymmetric expansion of PCs (Majda *et al.*, 2017; Sampathkumar *et al.*, 2014). Despite recent advances in the understanding of general principles underlying shape complexity, the molecular mechanisms and cellular networks that contribute to shape establishment are still largely elusive (Ivakov and Persson, 2013; Jacques *et al.*, 2014). Here, we provide experimental evidence that (i) positions IQD5 at cortical MT arrays in vegetative tissues (Fig. 1 and 2) and (ii) identifies IQD5 as a novel regulator of shape establishment in epidermal PCs of cotyledons and leaves (Fig. 3, 4 and 5). We further show that (iii) phenotypes in *iqd5* mutants correlate with alteration of cell wall properties (Fig. 6) and (iv) IQD5 recruits CaM to cortical MTs (Fig. 7). Thus, we provide first evidence for roles of Ca^2+^ signaling in the spatial coordination of cell expansion during interdigitated growth of PCs.

Our study revealed reduced lobe growth and growth restriction of neck regions in PCs of *iqd5* mutants. Lobe initiation on the other hand was only slightly reduced, and cell size as well as overall growth were unaffected, which indicates that IQD5 specifically functions in control of asymmetric expansion of PCs. Consistent with roles during asymmetric expansion, shape defects in *iqd5* mutants are established during early growth phases, i.e. in cotyledons at 2 DAG, at which lobe formation is initiated, and persist during later growth phases, in which cells expand within the lateral cell borders defined during the early growth phase (Zhang *et al.*, 2011). A role of IQD5 for regulating cellular expansion is further supported by its expression pattern identified in *pIQD5::GFP-GUS* reporter lines, which revealed uniform promoter activity within cotyledons and leaves. Similarly, growth and shape changes of PCs occur throughout the entire leaf, and growth rates display large heterogeneity between neighboring cells within expanding leaves (Elsner *et al.*, 2012). In contrast, cell cycle activity ceases in a longitudinal gradient during leaf maturation (Asl *et al.*, 2011). At later growth stages, cell division is restricted to the basal part of cotyledons and leaves, as indicated by analysis of the cell division marker CYCLINB1;1 (CYCB1;1) in transgenic *pCYC1;1::GUS* reporter lines (Carter *et al.*, 2017; Dhondt *et al.*, 2010; Ferreira *et al.*, 1994). The combined analysis of mutant phenotypes and of spatio-temporal expression domains thus establishes IQD5 as a novel factor controlling interdigitation of leaf epidermis PCs.

The development of wavy anticlinal cell walls from polyhedral precursor cells is proposed to rely on multi-polar growth patterns that involve subcellular variations in expansion rate (Fu *et al.*, 2005; Fu *et al.*, 2002). Asymmetric expansion and initiation of lobe formation correlate with local changes in cell wall composition and mechanical properties (Majda *et al.*, 2017), which are regulated by the plant cytoskeleton (Liu *et al.*, 2015). By analysis of GFP fluorescence in transgenic *pIQD5:IQD5-GFP/iqd5-1* lines, we demonstrate subcellular localization of IQD5-GFP to cortical MT arrays in leaf epidermal PCs. Functionality of the GFP-tagged IQD5 protein is indicated by efficient complementation of PC shape defects in *pIQD5:IQD5-GFP/iqd5-1* lines. MT localization was validated by oryzalin treatment, and is consistent with localization patterns of *pCaMV35S::YFP-IQD5* observed in transient expression assays in *N. benthamiana* leaves (Bürstenbinder *et al.*, 2017b). Quantification of fluorescence intensities along the outer periclinal cell wall suggests accumulation of IQD5-GFP at convex sides of indenting neck regions, which correlates with the preferential presence of parallel MT bundles at necks, as reported in several other studies (Armour *et al.*, 2015; Fu *et al.*, 2005; Sampathkumar *et al.*, 2014). Disturbance of MT organization, stability or dynamics by pharmacological agents or by mutations in MAPs, such as TRM2/LONGIFOLIA1 or KATANIN (KTN1), reduce cellular complexity of PC morphogenesis (Akita *et al.*, 2015; Lee *et al.*, 2006; Lin *et al.*, 2013). Most of the reported mutants with defects in PC shape, however, have pleiotropic effects, including reduced plant growth, organ twisting or swelling of cells (Qian *et al.*, 2009). Notably, although *IQD5* promoter activity was detectable in various vegetative tissues including roots and hypocotyls, *iqd5* mutants are macroscopically indistinguishable from the WT. Specific defects in PC morphogenesis thus indicate limited functional redundancy and compensation between the 33 IQD family members in Arabidopsis and point to unique roles of IQD5 in PC morphogenesis.

Discontiguous cell wall properties have been demonstrated to play an important role in cell shape establishment by introducing heterogeneity in mechanical properties of anticlinal walls (Majda *et al.*, 2017). Mechanical reinforcements in cell walls appear to be independent from local differences in cell wall thickness (Belteton *et al.*, 2018), but rather to rely on differences in cell wall composition (Majda *et al.*, 2017). Plants with impaired cellulose deposition, e.g. upon cellulase treatment or in mutants of the cellulose synthase AtCesA1 display strongly reduced lobing of PCs (Higaki *et al.*, 2016; Majda *et al.*, 2017). Direction of cellulose deposition is informed by the orientation of cortical MTs, which typically serve as tracks for PM localized cellulose synthase complexes (Endler and Persson, 2011). Histochemical staining of cellulose by calcofluor white revealed reduced deposition of cellulose in anticlinal walls of *iqd5* mutants. Thus, our data suggest that MT-localized IQD5 is required for efficient cellulose deposition, e.g., by controlling MT dynamics or organization, thereby changing subcellular sites of cellulose synthesis. Alternatively, IQD5 may mediate coupling of cellulose synthase movement to MT tracks, possibly by direct interaction with KINESIN LIGHT CHAIN-RELATED/CELLULOSE MICROTUBULE UNCOUPLING family members, which interact with Arabidopsis IQD1 at MTs (Bürstenbinder *et al.*, 2013; Liu *et al.*, 2016). Vice versa, cell wall mechanics also control MT orientation by executing subcellular and supracellular stresses (Sampathkumar *et al.*, 2014). Mechanical stress-induced rearrangements of MT arrays depend on KTN1 mediated MT severing, which occurs preferentially at crossover sites between MTs (Deinum *et al.*, 2017; Sampathkumar *et al.*, 2014). The severing activity of KTN1 may be affected by IQD5, e.g. by altering the degree of crossover events between individual MTs. To test these hypotheses, MT and CesA dynamics in PCs of *iqd5* mutants and the corresponding WT remain to be analyzed in future studies.

Intricate networks of cellular signaling pathways, which integrate developmental and environmental stimuli into cellular responses, coordinate the concerted action of the cell wall-PM-cytoskeleton continuum (Liu *et al.*, 2015). A central role in cellular signaling is played by the second messenger Ca^2+^, which regulates the MT cytoskeleton by yet unknown mechanisms (Hepler, 2016). The phenotypes in *iqd5* mutants, together with the IQD5-dependent recruitment of CaM to cortical MTs, provide first experimental evidence for functions of Ca^2+^ signaling in PC morphogenesis. Interestingly, Ca^2+^ signals are rapidly generated by exogenous application of auxin in several plant tissues, including hypocotyls, roots, and guard cells, and auxin induces rapid rearrangements in MT arrays (Vanneste and Friml, 2013). Lobing of PCs requires auxin-mediated activation of ROP signaling, which was proposed to depend on AUXIN BINDING PROTEIN1 (ABP1)-mediated auxin signaling and on polarized ditsributions of the PIN-FORMED1 (PIN1) auxin efflux carrier in lobe regions (Xu *et al.*, 2010). Recent reports, however, raise reasonable doubt about the validity of this model, as PC shape defects could be uncoupled from ABP1 functions in a novel *apb1* null mutant (Gao *et al.*, 2015) and polarized localization of PIN1 was not detectable in living cells of transgenic *pPIN1::PIN1-GFP* lines (Belteton *et al.*, 2018). Thus, despite experimental evidence for roles of auxin in lobe initiation, the mechanisms of action are largely elusive. IQD5 and related proteins of the IQD family may constitute promising candidates for integrating auxin signals into the reorganization of MT arrays, possibly via hormone-induced Ca^2+^ signals (Bürstenbinder *et al.*, 2017a; this study). Intriguingly, a recent study by Sugiyama *et al.* (2017) provides first indications for functions of IQD13 in spatial control of ROP signaling domains required for cell wall patterning during vessel development. A similar mechanism might apply to IQD5 during PC shape formation, thereby providing a potential link between auxin action, Ca^2+^ signaling and ROP GTPase activation. Collectively, our work identifies IQD5 as a novel regulator of PC shape and potential hub for coordination of cellular signaling, cytoskeletal reorganization, and cell wall remodeling. Our work thus provides a framework for future mechanistic studies of cellular signaling networks and of the cell wall-PM-MT continuum, which will aid a more holistic understanding of cellular processes guiding shape complexity.

## Acknowledgements

Funding was provided by the Deutsche Forschungsgemeinschaft (DFG, SFB648 to K.B. and S.A.), the German Academic Exchange Service (D.M.), the Erasmus Mundus program (P.K.), and by core funding of the Leibniz Association. This paper is a joint effort of the working group BIU, a unit of the German Centre for Integrative Biodiversity Research (iDiv) Halle-Jena-Leipzig, funded by the DFG (FZT 118).

## Supplementary data

Supplementary data are available at *JXB* online.

Fig. S1. *IQD5* expression analysis in *pIQD5*_*long*_*::GFP-GUS* reporter lines.

Fig. S2. Macroscopic analysis of growth parameters in the WT and *iqd5* mutants.

Fig. S3. Quantification and statistical analysis of PC shape features in 5-day-old seedlings of the WT and *iqd5* mutants.

Fig. S4. Quantification and statistical analysis of PC shape in cotyledons at 2 DAG.

Fig. S5. Quantification and statistical analysis of PC shape in cotyledons at 3 DAG.

Fig. S6. Quantification and statistical analysis of PC shape in cotyledons at 5 DAG.

Fig. S7. Quantification and statistical analysis of PC shape in cotyledons at 7 DAG.

Fig. S8. Quantification and statistical analysis of PC shape in cotyledons at 10 DAG.

Fig. S9. Quantification and statistical analysis of PC shape in true leaves.

